# Zero-determinant strategies under observation errors in repeated games

**DOI:** 10.1101/2020.01.17.910190

**Authors:** Azumi Mamiya, Genki Ichinose

## Abstract

Zero-determinant (ZD) strategies are a novel class of strategies in the repeated prisoner’s dilemma (RPD) game discovered by Press and Dyson. This strategy set enforces a linear payoff relationship between a focal player and the opponent regardless of the opponent’s strategy. In the RPD game, games with discounting and observation errors represent an important generalization, because they are better able to capture real life interactions which are often noisy. However, they have not been considered in the original discovery of ZD strategies. In some preceding studies, each of them has been considered independently. Here, we analytically study the strategies that enforce linear payoff relationships in the RPD game considering both a discount factor and observation errors. As a result, we first reveal that the payoffs of two players can be represented by the form of determinants as shown by Press and Dyson even with the two factors. Then, we search for all possible strategies that enforce linear payoff relationships and find that both ZD strategies and unconditional strategies are the only strategy sets to satisfy the condition. We also show that neither Extortion nor Generous strategies, which are subsets of ZD strategies, exist when there are errors. Finally, we numerically derive the threshold values above which the subsets of ZD strategies exist. These results contribute to a deep understanding of ZD strategies in society.

## I. INTRODUCTION

Cooperation is a basis for building sustainable societies. In a one-shot interaction, cooperation among individuals is suppressed because cooperation takes costs to the actor while defection does not. This cooperation-defection relationship is well captured by the prisoner’s dilemma (PD) game utilized in game theory. In the one-shot PD game, defection is the only Nash equilibrium. When the game is repeated, the situation drastically changes, which is modeled by the repeated prisoner’s dilemma (RPD) game [1]. In the RPD game, cooperation will be rewarded by the opponent in the future. In such a situation, cooperation becomes a possible equilibrium. This mechanism is called direct reciprocity [2–4] and makes it possible for players to mutually cooperate in the RPD game.

Evolutionary game theory (EGT) [5] studies how cooperation evolves in the RPD game. Among various cooperative strategies tested in evolutionary games, generous tit-for-tat [6] and win-stay lose-shift [7, 8] were robust to various kinds of evolutionary opponents under noisy conditions. EGT can find strong strategies against various opponents in evolving populations. One missing point was, what is a strong strategy against a direct opponent which utilizes any kind of strategy? In 2012, Press and Dyson suddenly answered this question from a different point of view. Using linear algebraic manipulations, they found a novel class of strategies which contain such ultimate strategies, called zero-determinant (ZD) strategies [9]. ZD strategies impose a linear relationship between the payoffs for a focal player and his opponent regardless of the strategy that the opponent implements. One of the subclasses of ZD strategies is Extortioner which never loses in a one-to-one competition in the RPD game against any opponents.

The discovery of ZD strategies stimulated many researchers. After Stewart and Plotkin raised a question [10], evolution or emergence of ZD strategies became one of the main targets in subsequent studies [11–25]. Then, this research spread in many directions including multiplayer games [19, 26–29], continuous action spaces [28–31], alternating games [31], asymmetric games [32], animal contests [33], human reactions to computerized ZD strategies [34, 35], and human-human experiments [28, 36, 37], which promote an understanding of the nature of human cooperation. For further understanding, see the recent elegant classification of strategies, partners (called “good strategies” in Ref. [11, 38]) and rivals, in direct reciprocity [39]. The utilization of ZD strategies has recently expanded to engineering fields, not just for human cooperation [40–42].

In those ZD studies, no errors were assumed. However, errors (or noise) are unavoidable in human interactions and they may lead to the collapse of cooperation due to negative effects. Thus, the effect of errors has been focused on in the RPD game [43–51]. However, only a few studies have concerned the effect of errors for ZD strategies [52, 53]. There are typically two types of errors: perception errors [45] and implementation errors [46]. Hao et al. [52] and Mamiya and Ichinose [53] considered the former case of the errors where players may misunderstand their opponent’s action because the players can only rely on their private monitoring [43, 47] instead of their opponent’s direct action. Those studies showed that ZD strategies can exist even in the case that such observation errors are incorporated. In those studies, no discount factor is considered. It is natural to assume that future payoffs will be discounted. Thus, some studies have focused on a discount factor for ZD strategies [30, 31, 54–56] and mathematically found the minimum discount factor above which the ZD strategies can exist [55].

In this study, we search for ZD strategies under the situations that observation errors and a discount factor are both incorporated. We search for the other possible strategies, not just ZD strategies, that enforce a linear payoff relationship between the two players. By formalizing the determinants for the expected payoffs in the RPD game, we mathematically found that only ZD strategies [9] and unconditional strategies [14, 55] are the two types which enforce a linear payoff relationship. We numerically show the threshold values above which the subsets of ZD strategies exist in the game.

## II. MODEL

### II.1. RPD with private monitoring

We consider the symmetric two-person repeated prisoner’s dilemma (RPD) game with private monitoring based on the literature [47, 52]. Each player *i* ∈ {*X*, *Y*} chooses an action *a_i_* ∈ {C, D} in each round, where C and D imply cooperation and defection, respectively. After the two players conduct the action, player *i* observes his own action *a_i_* and private signal *ω_i_* ∈ {*g*, *b*} about the opponent’s action, where *g* and *b* imply *good* and *bad*, respectively. In perfect monitoring, when the opponent takes the action C (D), the focal player always observes the signal *g* (*b*). In private monitoring, this is not always true. *σ*(***ω***|***a***) is the probability that a signal profile ***ω*** = (*ω_X_*, *ω_Y_*) is realized when the action profile is ***a*** = (*a_X_*, *a_Y_*) [47]. Let *ϵ* be the probability that an error occurs to one particular player but not to the other player while *ξ* be the probability that an error occurs to both players. Then, the probability that an error occurs to neither player is 1 – 2*ϵ* – *ξ*. For example, when both players take cooperation, *σ*((*g*, *g*)|(C, C)) = 1 – 2*ϵ* – *ξ*, *σ*((*g*, *b*)|(C, C)) = *σ*((*b*, *g*)|(C, C)) = *ϵ*, and *σ*((*b*, *b*)|(C, C)) = *ξ* are realized.

In each round, player *i*’s realized payoff *u_i_*(*a_i_*, *ω_i_*) is determined by his own action *a_i_* and signal *ω_i_*, such that *u_i_*(C, *g*) = *R*, *u_i_*(C, *b*) = *S*, *u_i_*(D, *g*) = *T*, and *u_i_*(D, *b*) = *P*. Note that the payoffs depend on the signals in private monitoring. Hence, his expected payoff is given by

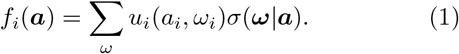

The expected payoff is determined by only action profile ***a*** regardless of signal profile ***ω***. Thus, the expected payoff matrix is given by

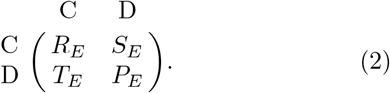

According to Eq. (1), *R_E_*, *S_E_*, *T_E_*, and *P_E_* are derived as *R_E_* = *R*(1 – *ϵ* – *ξ*) + *S*(*ϵ* + *ξ*), *S_E_* = *S*(1 – *ϵ* – *ξ*) + *R*(*ϵ* + *ξ*), *T_E_* = *T*(1 – *ϵ* – *ξ*) + *P*(*ϵ* + *ξ*), *P_E_* = *P*(1 – *ϵ* – *ξ*) + *T*(*ϵ* + *ξ*), respectively. We assume that

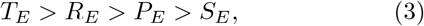

and

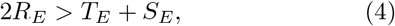

which dictate the RPD condition with observation errors.

In this paper, we introduce a discount factor to the RPD game with private monitoring. The game is to be played repeatedly over an infinite time horizon but the payoff will be discounted over rounds. Player *i*’s discounted payoff of action profiles ***a***(*t*), *t* ∈ {0, 1, …, ∞} is *δ^t^f_i_*(***a***(*t*)) where *δ* is a discount factor and *t* is a round. This game can be interpreted as repeated games with a finite but undetermined time horizon. Finally, the average discounted payoff of player *i* is

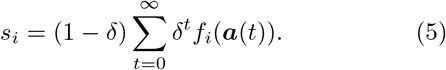

### II.2. Determinant form of expected payoff in the RPG game

Here, we proceed to show that Eq. (5) can be represented by a determinant form even for the repeated games with observation errors and a discount factor, as Press and Dyson did for the repeated game without error and no discount factor [9]. The action profiles ***a***(*t*) in Eq. (5) need to be specified to calculate *s_i_*. Those profiles are determined after the strategies of two players are given. Consider player *i* that adopts memory-one strategies, with which they can use only the outcomes of the last round to decide the action to be submitted in the current round. A memory-one strategy is specified by a 5-tuple; *X*’s strategy is given by a combination of

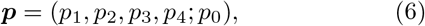

where 0 ≤ *p_j_* ≤ 1, *j* ∈ {0, 1, 2, 3, 4}. The subscripts 1, 2, 3, and 4 of *p* mean previous outcomes C*g*, C*b*, D*g*, and D*b*, respectively. In Eq. (6), *p*_1_ is the conditional probability that *X* cooperates when *X* cooperated and observed signal *g* in the last round, *p*_2_ is the conditional probability that *X* cooperates when *X* cooperated and observed signal *b* in the last round, *p*_3_ is the conditional probability that *X* cooperates when *X* defected and observed signal *g* in the last round, and *p*_4_ is the conditional probability that *X* cooperates when *X* defected and observed signal *b* in the last round. Finally, *p*_0_ is the probability that *X* cooperates in the first round. Similarly, *Y*’s strategy is specified by a combination of

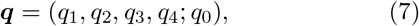

where 0 ≤ *q_j_* ≤ 1, *j* ∈ {0, 1, 2, 3, 4}.

Define ***v***(*t*) = (*v*_1_(*t*), *v*_2_(*t*), *v*_3_(*t*), *v*_4_(*t*)) as the stochastic state of two players in round *t* where the subscripts 1, 2, 3, and 4 of *v* imply the stochastic states (C,C), (C,D), (D,C), and (D,D), respectively. *v*_1_ (*t*) is the probability that both players cooperate in round *t*, *v*_2_(*t*) is the probability that *X* cooperates and *Y* defects in round *t*, and so forth. Then, the expected payoff to player *X* in round *t* is given by ***v***(*t*)***S**_X_*, where 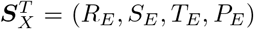. The expected per-round payoff to player *X* in the repeated game is given by

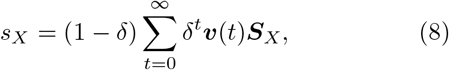

where 0 < *δ* < 1. The initial stochastic state is given by

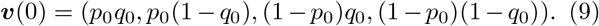

The state transition matrix *M* of these repeated games with observation errors is given by

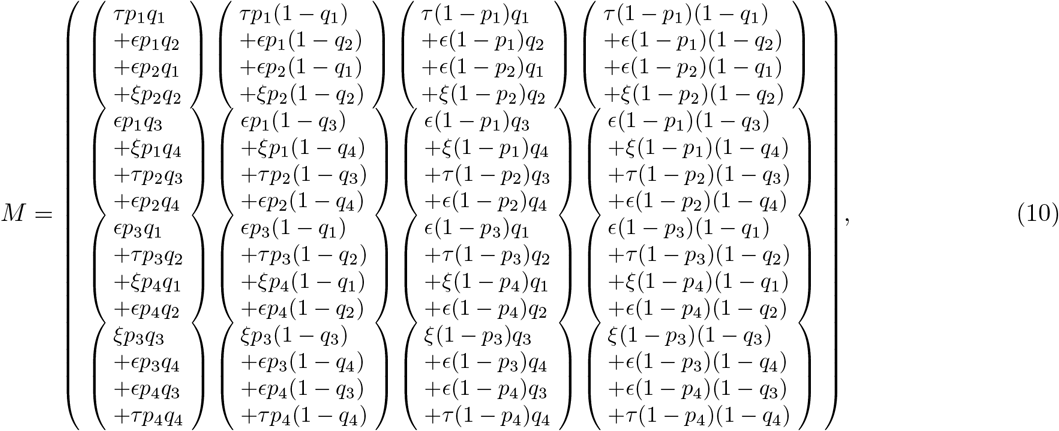

where *τ* =1 – 2*ϵ* – *ξ*. Then, we obtain

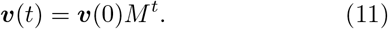

By substituting Eq. (11) in Eq (8), we obtain

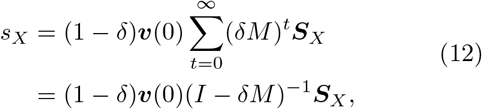

where *I* is the 4 × 4 identity matrix. Then, let

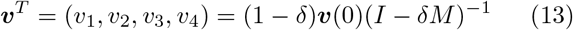

be the mean distribution of ***v***(*t*). Additionally, we define

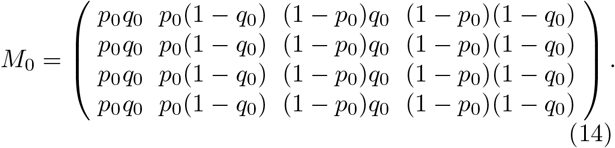

Because *v*_1_ + *v*_2_ + *v*_3_ + *v*_4_ = 1 (Appendix A), the following holds (Appendix B):

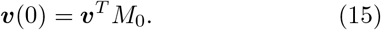

By substituting Eq. (15) in Eq. (13) and multiplying both sides of the equation by (*I* – *δM*) from the right, we obtain

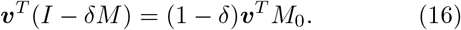

Equation (16) and *M*′ ≡ *δM* + (1 – *δ*)*M*_0_ – *I* yield

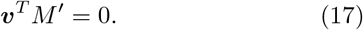

With Eq. (17), we immediately obtain a formula for the dot product of an arbitrary vector ***f**^T^* = (*f*_1_, *f*_2_, *f*_3_, *f*_4_) with the fourth column vector ***u*** of matrix *M*′ as a consequence of Press and Dyson’s formalism, which can be represented by the form of the determinant

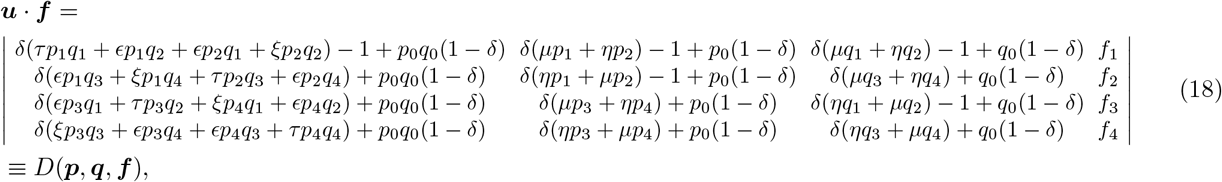

where *μ* = 1 – *ϵ* – *ξ* and *η* = *ϵ* + *ξ*. Furthermore, Eq. (18) should be normalized to have its components sum to 1 by ***u*** · **1**, where **1** = (1, 1, 1, 1). Then, we obtain the dot product of an arbitrary vector ***f*** with mean distribution ***v***. Replacing the last column of *D*(***p***, ***q***, ***f***) with player *X*’s and *Y*’s expected payoff vector, respectively, we obtain their per-round expected payoffs:

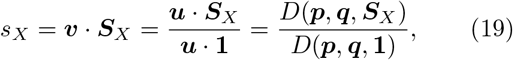

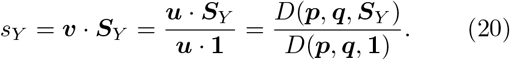

When we set *δ* =1, Eq. (18) corresponds to Eq. (2) of [52]. By using Eq. (18), we can calculate players’ per-round expected payoffs when 0 < *δ* ≤ 1 by the form of the determinants. *δ* = 1 is the case where future payoffs are not discounted.

## III. RESULTS

### III.1. All strategies that enforce linear payoff relationships

Since we are interested in the payoff relationship between the two players, we linearly combine those payoffs represented by Eqs. (19) and (20). The linear combination of *s_X_* and *s_Y_* can also be represented by the form of the determinant:

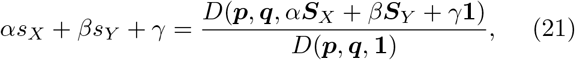

where *α, β*, and *γ*, are arbitrary constants. The numerator of the right side of Eq. (21) is expressed in the following way:

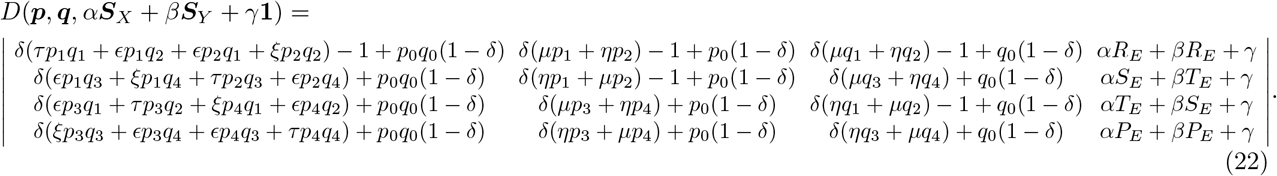

If Eq. (22) is zero, the relationship between the two players’ payoffs becomes linear:

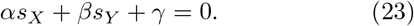

Thus, we search for all of the solutions such that *D*(***p**, **q**, α**S**_X_* + *β**S**_Y_* + *γ***1**) = 0.

Press and Dyson [9] (without error) and Hao et al. [52] (with observation errors) searched for the case that second and fourth columns of the determinant take the same value. This makes the determinant zero. Also, Mamiya and Ichinose [53] searched for all the cases, from all possibilities, that make the determinant zero with observation errors. Here, we extend Mamiya and Ichinose [53] to the case with both observation errors and a discount factor.

The following determinant theorem gives such a condition:

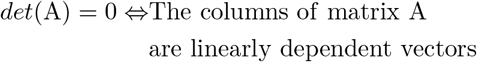

for *n* × *n* matrix A. We define ***d**_i_*, *i* ∈ {1, 2, 3, 4} as *i*-th column vector of the determinant of Eq. (22). From the above theorem, if the columns of the determinant of Eq. (22) are linearly dependent vectors, there exist real numbers *s*, *t*, *u*, *v*, *α*, *β*, and *γ*, except for the trivial solution ((*s*, *t*, *u*, *v*) = (0, 0, 0, 0),(*α*, *β*, *γ*) = (0, 0, 0)), such that

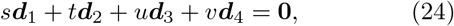

where vector **0** denotes a zero vector. We give the detailed calculation in Appendix C.

As a result, we found that, in the RPD game even with observation errors (imperfect monitoring) and a discount factor, the only strategies that impose a linear payoff relationship between the two players’ payoffs are either

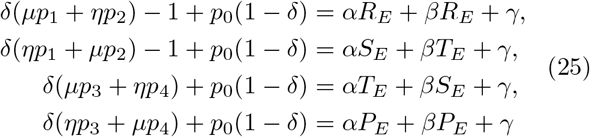

or

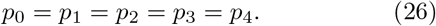

The former corresponds to ZD strategies and the latter corresponds to unconditional strategies, respectively.

### III.2. Extortion and Generous no longer exist when there are errors

Extortion [9] and Generous [23] strategies are well-known subsets of ZD strategies, which have important characteristics. Extortion never loses to any opponent in a one-to-one competition in terms of the expected payoffs. Moreover, they finally impose ALLC (always cooperate) to the opponent who tries to improve his payoff [9]. On the other hand, Generous strategies always obtain lower payoffs than the opponent except for mutual cooperation. Hence, Generous strategies are known as one of the cooperative ZD strategies. Generous strategies are weak in a one-to-one competition. However, in a large evolving population, cooperative groups are more successful than the group of Extortioners. Thus, evolution leads from Extortion to Generous strategies.

In Eq. (25), we substitute *α* = *ϕ*, *β* = – *ϕχ*, and *γ* = *ϕ*(*χ* – 1)*κ* [55] to obtain

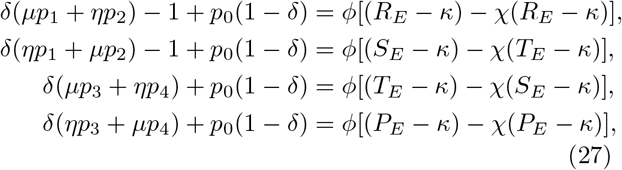

where *κ* = *P_E_* with 1 ≤ *χ* < ∞ represents Extortion while *κ* = *R_E_* with 1 ≤ *χ* < ∞ represents Generous strategies. The parameter *χ* gives the correlation between two players’ payoffs. Thus, we call *χ* a correlation factor. The parameter *κ* corresponds to the payoff that the ZD strategy would obtain against itself. We thus call *κ* baseline payoff as used in Ref. [14].

Those two prominent strategy sets exist when there are no errors. Here, by contrast, we prove that Extortion and Generous strategies no longer exist when there are errors. The detail calculation is provided in Appendix D.

However, Hao et al. [52] and Mamiya and Ichinose [53] found that there exist ZD strategies which partially have the characteristic of Extortion even when there are errors. In the original meaning, Extortion has two properties: (1) his payoff increase is always higher than any opponent’s due to *χ* > 1 (payoff control ability) and (2) he is not outperformed by anyone (payoff dominance). When there are errors, the second property is lost although the first property remains. Intuitively, this is because errors introduce uncertainty into the payoffs and, consequently, enforce a negative impact on the accuracy of player *X*’s payoff-based strategy setting. To keep the payoff control ability, the payoff dominance needs to be sacrificed when there are errors. Hao et al. called the ZD strategies which only have the first property *contingent extortion* [52].

### III.3. Existence of subsets of ZD strategies

Since observation errors and a discount factor are considered, in general, the ranges in which ZD strategies can exist are narrowed. Ichinose and Masuda mathematically showed the minimum threshold values above which Equalizer (another subclass of ZD strategies), Extortion, and Generous strategies can exist [55]. Here, we numerically address threshold values where subsets of ZD strategies can exist.

#### Minimum discount factor for Equalizer

Equalizer strategies are a subclass of ZD strategies. They can fix the expected payoffs of the opponent regardless of the opponent’s strategies [9]. We first show minimum discount factor *δ_c_* for Equalizer when observation errors *ϵ* and *ξ* are given. Equalizer can fix the opponent payoff no matter what the opponent takes, which means that

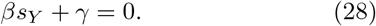

This is obtained by substituting *α* = 0 into Eq. (23). Note that *χ* → ∞ in Eq. (27) corresponds to Equalizer [52]. We substitute *α* = 0 into Eq. (25) to obtain Equalizer:

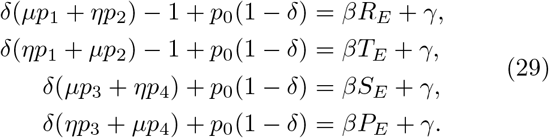

If we solve Eq. (29) for *β, γ, p*_2_ and *p*_3_,

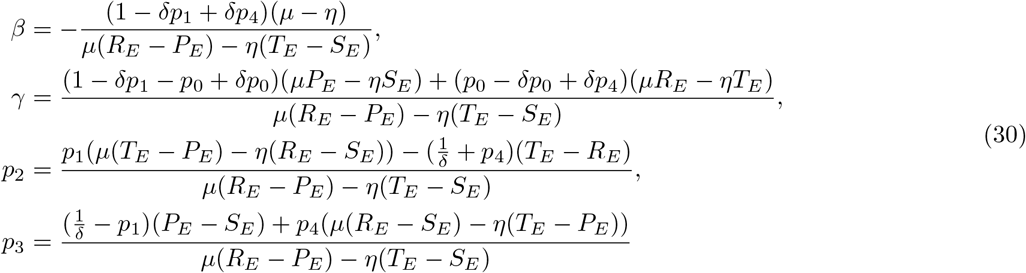

are obtained. By substituting *β* and *γ* into Eq. (28), player *Y*’s payoff is fixed at

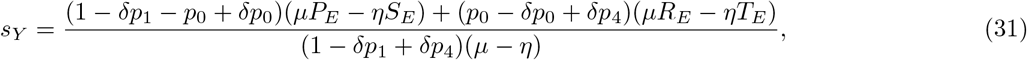

which is independent of the opponent’s strategies ***q***. Equations (30) and (31) correspond to Eq. (10) in [52] when *δ* = 1.

It should be noted that Equalizer does not require any condition on *p*_0_. However, Eq. (31) indicates that the payoff that Equalizer enforces on the opponent’s payoff *s_Y_* depends on the value of *p*_0_. Because Eq. (31) is a weighted average of (*μP_E_* – *ηS_E_*)/(*μ* – *η*) and (*μR_E_* – *ηT_E_*)/(*μ* – *η*) with non-negative weights, Equalizer can impose any payoff value *s_Y_* such that (*μP_E_* – *ηS_E_*)/(*μ* – *η*) ≤ *s_Y_* ≤ (*μR_E_* – *ηT_E_*)/(*μ* – *η*) as long as error rate *η* = *ϵ* + *ξ* is not so high. If (*μP_E_* – *ηS_E_*)/(*μ* – *η*) is enforced, it holds that *p*_0_ – *δp*_0_ + *δp*_4_ = 0, and hence *p*_4_ = *p*_0_ = 0. If (*μR_E_* – *ηT_E_*)/(*μ* – *η*) is enforced, it holds that 1 – *δp*_1_ – *p*_0_ + *δp*_0_ = 0, and hence *p*_1_ = *p*_0_ = 1.

Equalizer must satisfy the condition 0 ≤ *p_i_* ≤ 1 in Eq. (29). The existence of Equalizer strategies also depends on *δ*, *ϵ*, and *ξ*. We numerically find the minimum discount factor *δ_c_* and the condition of (*ϵ*, *ξ*) for which Equalizer exists. *δ* ≥ *δ_c_* is the condition for *δ* under which Equalizer strategies exist. Figure 1A shows *δ_c_* when *ϵ* + *ξ* is given. We set (*T, R, P, S*) = (1.5, 1, 0, −0.5) and excluded the case *ϵ*+*ξ* > 1/3 because *T_E_* > *R_E_* > *P_E_* > *S_E_* is not satisfied under the situation. Note that the effects of *ϵ* and *ξ* are the same because *η* = *ϵ*+*ξ* and *μ* = 1 – *ϵ* – *ξ* in Eq. (29) includes both *ϵ* and *ξ*. When there was no error (*ϵ* + *ξ* = 0), *δ_c_* was about 0.33. When the errors were *ϵ* + *ξ* = 0.1 and 0.2, *δ_c_* were about 0.52 and 0.93. As a result, we found that *δ* ≥ *δ_c_* for Equalizer becomes larger as the error is increased. Figure 1B shows the possible payoff range that Equalizer can enforce to the opponent (*s_Y_*) when corresponding *ϵ* + *ξ* is given. If there is no error (*ϵ* + *ξ* = 0), Equalizer can enforce all possible payoffs between *P_E_* = 0 and *R_E_* = 1 as shown in Ichinose and Masuda [55]. However, as error rates become larger, the possible range of payoffs that Equalizer can enforce becomes smaller. When the errors were *ϵ* + *ξ* = 0.1 and 0.2, the possible payoff ranges were 0.21 ≤ *s_Y_* ≤ 0.79 and 0.47 ≤ *s_Y_* ≤ 0.53, respectively.

**FIG. 1.**
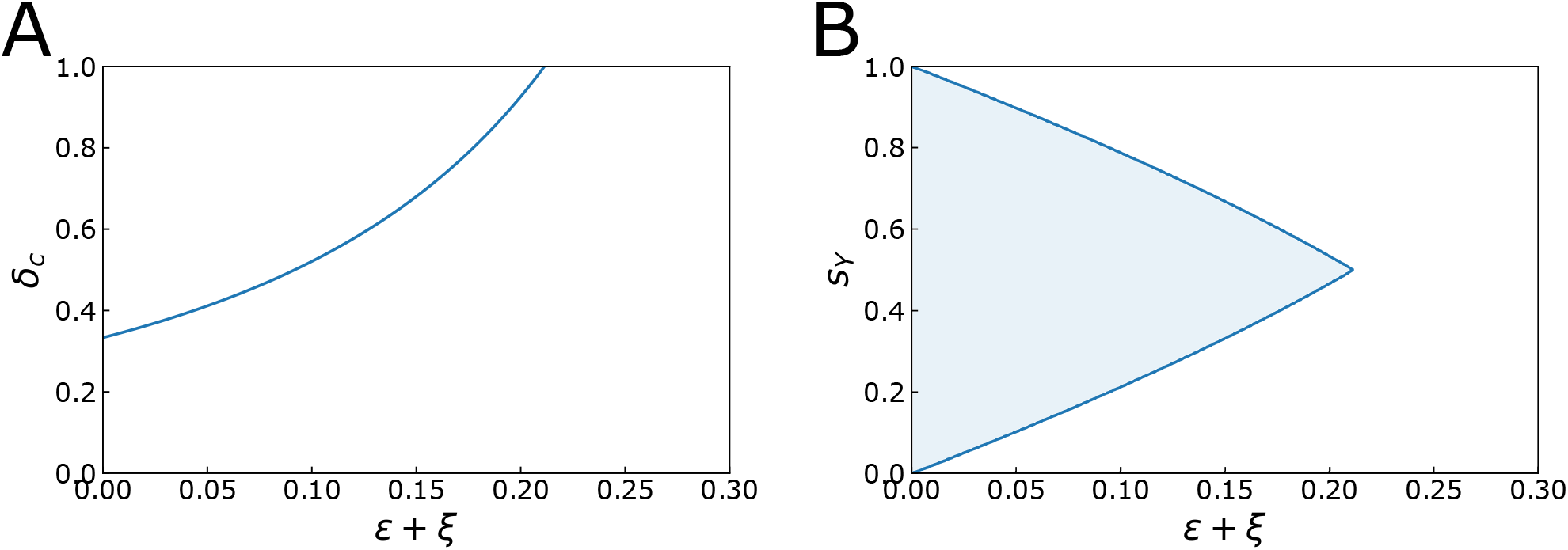
(A) Minimum discount factor *δ_c_* for Equalizer. (B) The possible payoff range that Equalizer can enforce to the opponent (*s_Y_*) when corresponding *ϵ* + *ξ* is given.

#### Minimum correlation factor for ZD strategies with 1 ≤ *χ* < ∞

We numerically calculated the minimum correlation factor *χ_c_* for subsets of ZD strategies with 1 ≤ *χ* < ∞ to exist (Fig. 2). Each curve corresponds to each *δ* as shown in the legend. The area surrounded by each curve and the vertical axis is the region of *χ* which can be utilized by the ZD strategies when *ϵ*+*ξ* is fixed. As the error *ϵ*+*ξ* becomes larger and the discount factor *δ* becomes smaller, the minimum correlation factor *χ_c_* becomes larger.

**FIG. 2.**
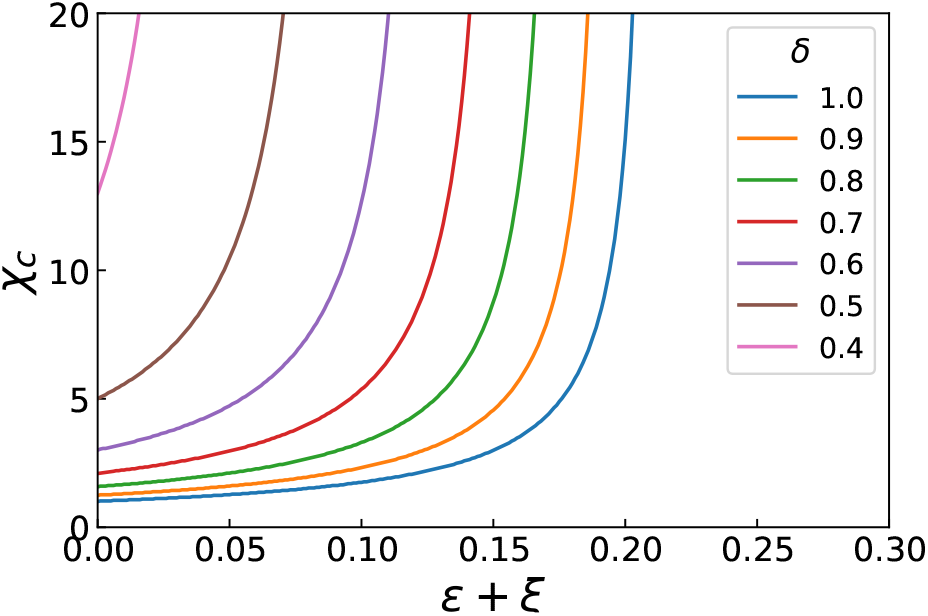
Minimum correlation factor *χ_c_* for subsets of ZD strategies with 1 ≤ *χ* < ∞. (*T, R, P, S*) = (1.5, 1, 0, −0.5). We adopted *p*_0_ and *κ* so that *χ_c_* was minimized.

### III.4. Numerical examples of representative ZD and unconditional strategies under errors in repeated games

We numerically demonstrate that ZD and unconditional strategies can impose a linear relationship between the two players’ payoffs while others cannot in the RPD game under errors. We take up contingent Extortion and Equalizer as the representative of ZD strategies, ALLD (always defect) as the representative of unconditional strategies, and Win-Stay-Lose-Shift (WSLS) as neither ZD nor unconditional strategies in general.

Figure 3 shows the relationship between the two players’ expected payoffs per game with payoff vector (*T, R, P, S*) = (1.5, 1, 0, −0.5). The gray quadrangle in each panel represents the feasible set of payoffs. We fixed one particular strategy for player *X* (vertical line) and randomly generated 1,000 strategies that satisfy 0 ≤ *q*_0_, *q*_1_, *q*_2_, *q*_3_, *q*_4_ ≤ 1 for player *Y* (horizontal axis). Thus, each black dot represents the payoff relationship between two players. In addition, the blue and red are the particular cases for player *Y*. Red is the case that player *Y* is ALLD and blue is the case that player *Y* is ALLC. We set *δ* =1 for Figs. 3A–D and *δ* = 0.9 for Figs. 3E–H. In each figure, we used three error rates *ϵ* + *ξ* = 0, 0.1, and 0.2.

**FIG. 3.**
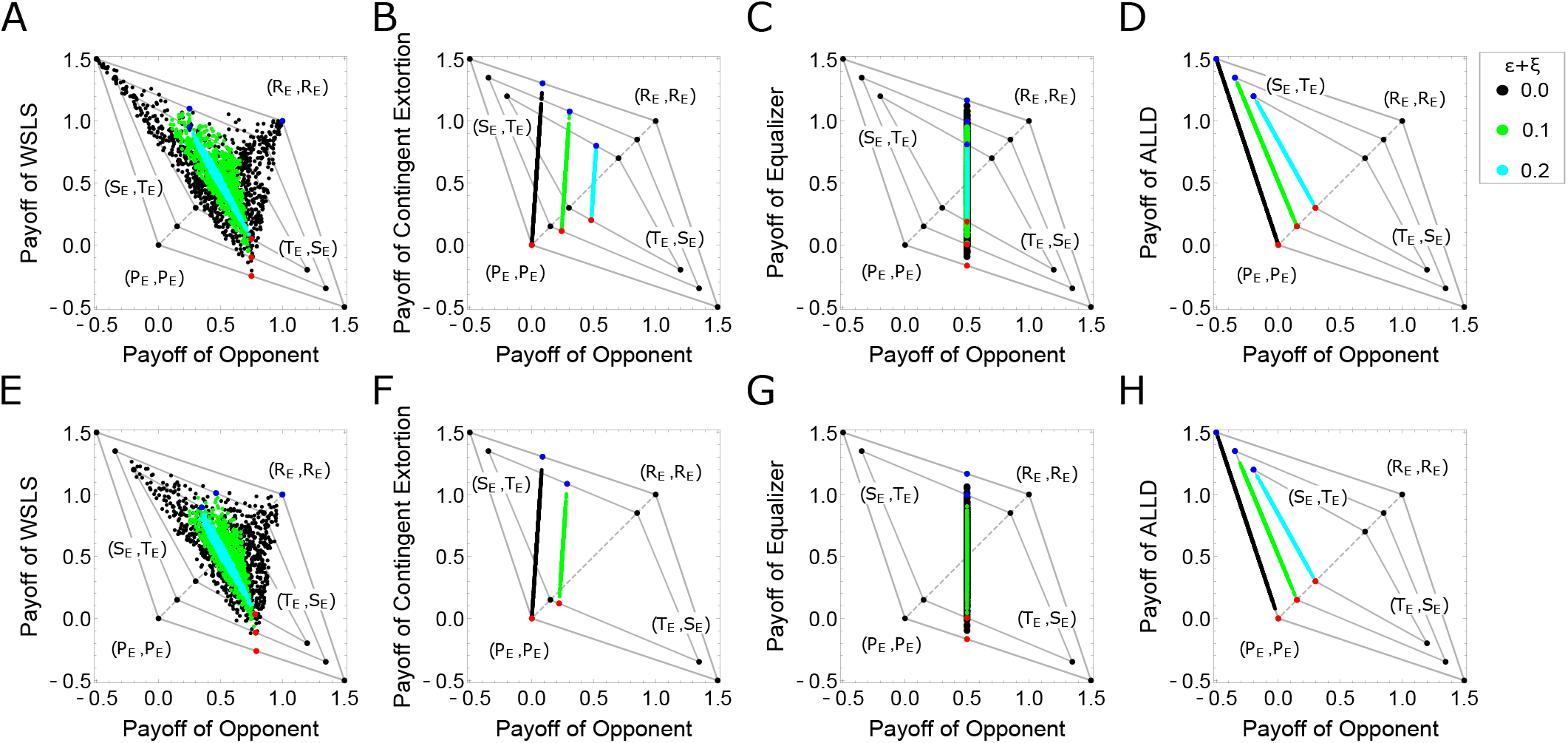
The payoff relationships between two players in the RPD game under observation errors. Payoff vector: (*T, R, P, S*) = (1.5, 1, 0, −0.5). (A,E) WSLS strategy vs. 1000 + 2 strategies. (B,F) Contingent Extortioner strategy vs. 1000 + 2 strategies. (C,G) Equalizer strategy vs 1000 + 2 strategies. (D,H) ALLD strategy vs. 1000 + 2 strategies. (A)–(D) are the case of *δ* = 1 (no discount factor), and (E)–(H) correspond to (A)–(D) when *δ* = 0.9, respectively.

Figures 3A and E show the case with a WSLS strategy vs. 1000 + 2 strategies. In that case, *ξ* = 0 is fixed and *ϵ* is caried to 0, 0.1, 0.2. As WSLS strategies are neither ZD nor unconditional strategies in general, the payoff relationships are not linear irrespective of errors and a discount factor.

Figures 3B and F show the case with an contingent Extortion vs. 1000 + 2 strategies. If there are no errors, Extortion is unbeatable against any opponent as shown by black dots. For instance, when *δ* = 1 and *ϵ* + *ξ* = 0, Extortioner ***p*** = (0.86, 0.77, 0.09, 0) which passes over (*P_E_, P_E_*) can impose a linear payoff relationship to the opponent, with the slope *χ* = 15 [black dots in Fig. 3B]. Note that *δ* =1 corresponds to a no discounting game. In that case, the expected payoff of a game does not depend on *p*_0_ because all terms which contain *p*_0_ and *q*_0_ in Eq. (18) vanish. In other words, the value of *p*_0_ is arbitrary. Even if *δ* = 0.9 and *ϵ* + *ξ* = 0, Extortioner ***p*** = (0.955556, 0.855556, 0.1, 0; 0) which passes over (*P_E_*, *P_E_*) can impose a linear payoff relationship to the opponent, with the slope *χ* = 15 [black dots in Fig. 3F].

However, as proved in Sec. III.2, when there are errors, only contingent Extortion can exist. Thus, there exists the region that the expected payoff of the contingent Extortion is lower than the opponent’s payoff near (*P_E_, P_E_*) [see yellow-green and cyan dots in Figs. 3B and F] even though the increase of the strategies is still larger than the opponent’s due to *χ* > 1 when the opponent tries to increase his payoff.

When *δ* = 1 and *ϵ* + *ξ* = 0.1, contingent Extortioner ***p*** = (0.926875, 0.818125, 0.111875, 003125) which passes over (*P_E_* + 0.1, *P_E_* + 0.1) can impose a linear payoff relationship to the opponent, with the slope *χ* = 15 [yellow-green dots in Fig. 3B]. Even if *δ* = 0.9 and *ϵ* + *ξ* = 0.1, contingent Extortioner ***p*** = (0.941667, 0.7, 0.241667, 0; 0) which has the same slope *χ* = 15 can still exist [yellowgreen dots in Fig. 3F]. Nevertheless, these two contingent Extortioners’ expected payoffs are lower than the opponents near (*P_E_, P_E_*). When *δ* = 1 and *ϵ* + *ξ* = 0.2, contingent Extortioner ***p*** = (1, 0.86, 0.14, 0) which passes over (*P_E_* + 0.2, *P_E_* + 0.2) can impose a linear payoff relationship to the opponent, with the slope *χ* = 15 [cyan dots in Fig. 3B]. However, this contingent Extortioner’s payoff is lower than the opponent’s near (*P_E_*, *P_E_*), too. When *δ* = 0.9 and *ϵ* + *ξ* = 0.2, there is no contingent Extortioner as shown in Fig. 2.

Figures 3C and G show the case with an Equalizer strategy vs. 1000 + 2 strategies. When *δ* = 1 and *ϵ* + *ξ* = 0, 0.1, and 0.2, Equalizers ***p*** = (2/3, 1/3, 2/3, 1/3), ***p*** = (0.8, 0.365217, 0.634783, 0.2), and ***p*** = (0.99, 0.74, 0.26, 0.01) can fix the opponents’ (player *Y*) expected payoffs at *s_Y_* = 0.5 irrespective of *Y*’s strategies, respectively [black, yellow-green, and cyan dots in Fig. 3C]. Also, when *δ* = 0.9 and *ϵ*+*ξ* = 0 and 0.1, Equalizer ***p*** = (2/3, 0.277778, 0.722222, 1/3; 1/2), and ***p*** = (0.833333, 0.350242, 0.649758, 1/6; 1/2) can fix the opponents’ (player *Y*) expected payoffs at *s_Y_* = 0.5 irrespective of *Y*’s strategies, respectively [black and yellow-green dots in Fig. 3G]. When *δ* = 0.9 and *ϵ* + *ξ* = 0.2, there exists no Equalizer as shown in Fig. 1.

Lastly, we show the case of ALLD [Figs. 3D and H]. ALLD strategy is one of the unconditional strategies where we set *r* = 0 in ***p*** = (*r, r, r, r; r*), 0 ≤ *r* ≤ 1. As shown in Eq. (C15) and Figs. 3D and H, *δ* does not affect the expected payoff between both players. When *ϵ* + *ξ* = 0, 0.1, and 0.2, those linear equations are *s_X_* + 3*s_Y_* = 0 (black), *s_X_* + 2.4*s_Y_* – 0.51 = 0 (yellow-green), and *s_X_* + 1.8*s_Y_* – 0.84 = 0 (cyan), respectively [53].

## IV. DISCUSSION

We considered both a discount factor and observation errors in the RPD game and analytically studied the strategies that enforce linear payoff relationships in the game. First, we successfully derived the determinant form of the two players’ expected payoffs even though a discount factor and observation errors are incorporated. Then, we searched for all possible strategies that enforce linear payoff relationships in the RPD game. As a result, we found that both ZD strategies and unconditional strategies are the only strategy sets to enforce the relationship to the opponent. Then, we proved that Extortion and Generous strategies no longer exist when there are errors. Finally, we numerically showed minimum discount factors for Equalizer (*χ* → ∞) and minimum correlation factors for other subsets of ZD strategies (1 ≤ *χ* < ∞) above which those ZD strategies exist.

We showed that ZD strategies can still exist even when discount factor *δ* deviates from 1. In real life, we remember interacting with other people but the interaction will conclude after a certain time. People sometimes change who they interact with because they move to other places. Young animals spend most of their time with their parents. After animals grow up and become adults, they interact with new companions. There are interactions in a short time such as mating. Our results demonstrate that strategies which unilaterally control the opponents’ payoffs exist even in those limited repeated interactions.

We also showed that ZD strategies can still exist even when there are observation errors to some extent. Such noise often happens in real interactions between individuals. Thus, if ZD strategies were to exist only without noise, the applicability of ZD strategies in real problems would be quite limited. Our results are important because we expanded the applicability of ZD strategies to those problems.

Our results are limited to the two-player RPD games. Other studies have focused on *n*-player games [19, 26, 27, 56]. It is worth investigating games including observation errors and a discount factor for *n*-player games. On the other hand, regarding memory, our study only used memory-1 strategies. A recent study revealed the role of longer memories for the evolution of cooperation, which is another direction to investigate [51].

When spatial structures are included, the different role of Extortioner has been known [16–18, 21]. Extortioners are neutral with respect to ALLDs if there are no errors. Thus, Extortioners can neutrally invade the sea of ALLDs in a spatial structure. On the other hand, the best response to Extortioners is ALLC. Once ALLC happens, the clusters of ALLC are better than those of the Extortioner. Then, cooperation is promoted. In this way, it has been demonstrated that Extortion acts as a catalyst for cooperation. Another interest is how observation errors and a discount factor affect the evolution of cooperation in a spatial setting.

We considered a model of private monitoring in direct reciprocity where signals are personally interpreted. Signals would play an important role in indirect reciprocity rather than direct reciprocity because signals are shared by many players as a social norm which affects their behaviors. As far as we know, no one has found ZD strategies in indirect reciprocity. We do not even know that we can use similar techniques for the problem. Thus, this is a potentially challenging and exciting direction of future research.

Game theory has been applied to practical problems. For instance, it has been used in the problem of selfish routing or traffic congestion. In those problems, imposing a tax on selfish terminals or drivers is one possible strategy to improve the efficiency of the total flow. As we showed, ZD strategies can induce cooperative behavior to the other players. Thus, incorporating ZD strategies instead of imposing taxes in those problems may be an effective way to improve efficiency. Our study contributes to new research directions of ZD strategies to various fields.

## Appendix A: Proof of *v*_1_ + *v*_2_ + *v*_3_ + *v*_4_ = 1

We show that the sum of elements in the mean distribution ***v*** = (*v*_1_, *v*_2_, *v*_3_, *v*_4_) is equal to 1. We define

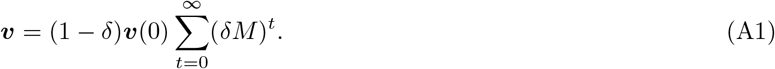

This is another form of Eq (13). Because the sum of every row in the transition matrix *M* is equal to one, the sum of every row of 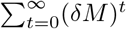 is equal to 1/(1 – *δ*). The sum of vector elements in ***v***(0) is unchanged from 1 even if the vector is multiplied by 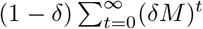. Therefore, *v*_1_ + *v*_2_ + *v*_3_ + *v*_4_ = 1 holds.

## Appendix B: Calculation of *v*(0) = *v^T^ M*_0_

We show that ***v***(0) and ***v**^T^M*_0_ are equal. ***v*** and *M*_0_ are defined by Eqs. (13) and (14), respectively. We calculate the matrix multiplication ***v**^T^M*_0_.:

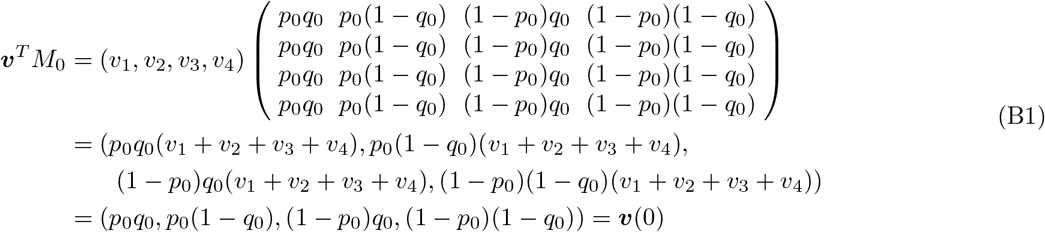

Therefore, the following holds:

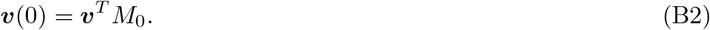

## Appendix C: Strategies that enforce *D*(*p*, *q*, *αS_X_* + *βS_Y_* + *γ*1) = 0

To search for all possible strategies that make *D*(***p***, ***q***, *α**S**_X_* + *β**S**_Y_* + *γ***1**) = 0, we express Eq. (24) in component form:

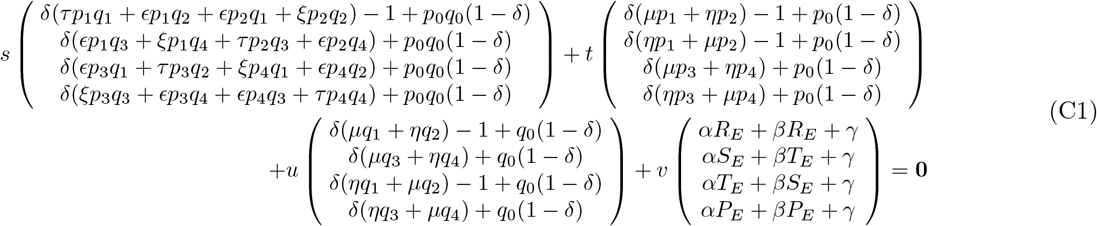

By taking out ***q*** from Eq. (C1), we obtain

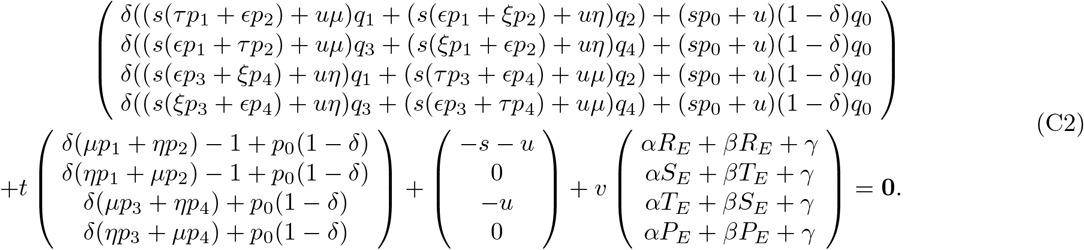

Here, we search for strategies which satisfy *D*(***p***, ***q***, *α**S**_X_* + *β**S**_Y_* + *γ***1**) = 0 irrespective of *Y*’s strategy ***q***, meaning that Eq. (C2) must hold true irrespective of ***q***. Therefore, the coefficients of each element ***q*** in Eq. (C2) must equal zero, that is, the following conditions are necessary:

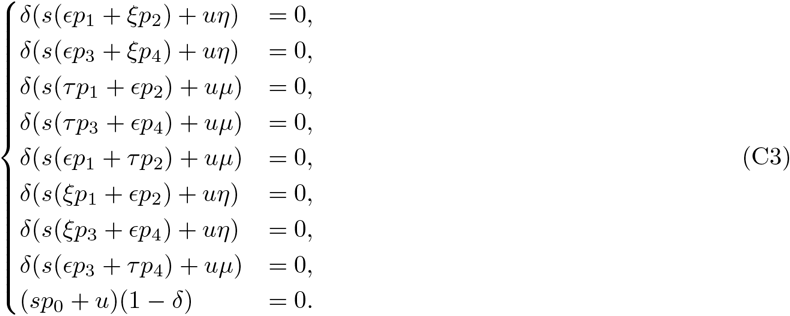

When Eq. (C3) holds, the first terms of Eq. (C2) are eliminated and we obtain

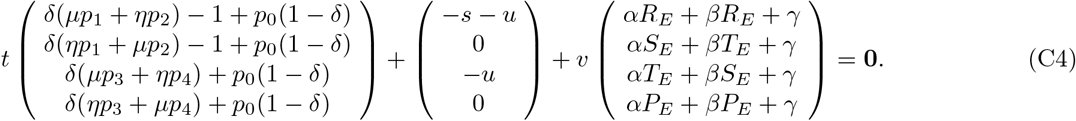

If there exist real numbers, *s, t, u, v, α, β*, and *γ* such that Eqs. (C3) and (C4) are satisfied simultaneously, *D*(***p***, ***q***, *α**S**_X_* + *β**S**_Y_* + *γ***1**) = 0 holds irrespective of ***q***. We first solve Eq. (C3). After some calculations, Eq. (C3) becomes

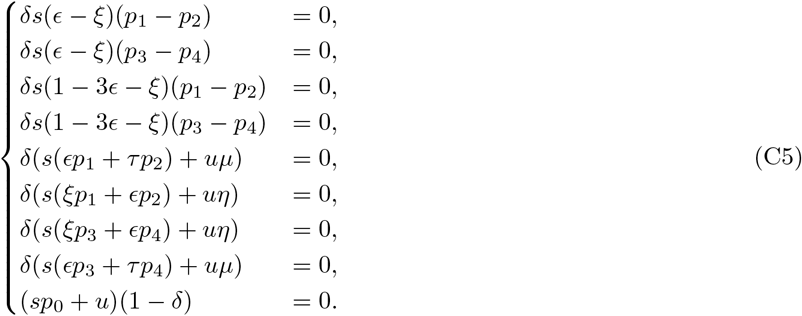

When we solve the first four equations, we obtain (1) *s* = 0, (2) *ϵ* – *ξ* = 0 and 1 – 3*ϵ* – *ξ* = 0, (3) *p*_1_ – *p*_2_ =0 and *p*_3_ – *p*_4_ = 0. We further analyze whether these solutions satisfy the last four equations and Eq. (C4) by dividing them into four cases as follows.

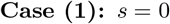

In this case, we substitute *s* = 0 into Eq. (C5) to obtain

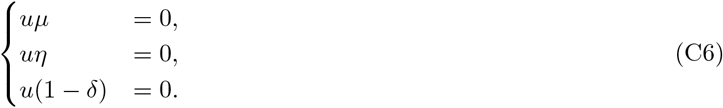

The equations *μ* = 0 and *η* = 0 do not hold at the same time due to *μ* = 1 – *ϵ* – *ξ* and *η* = *ϵ* + *ξ*. Therefore, one of the solutions of Eq. (C5) is *s* = 0 and *u* = 0. Next, we check whether this solution satisfies Eq. (C4). We substitute *s* = 0 and *u* = 0 into Eq. (C4) to obtain

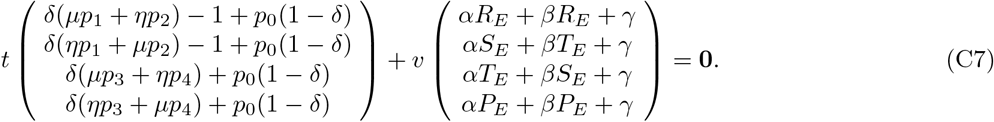

Here, when we set *t* = 0, either equation

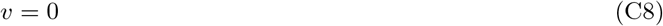

or

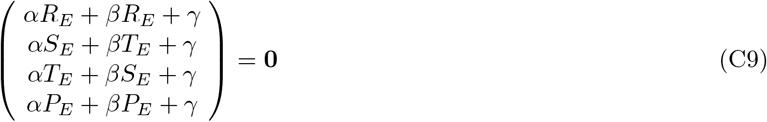

must hold. When we set *v* = 0, we obtain the trivial solution (*s, t, u, v*) = (0, 0, 0, 0). Also, when we solve Eq. (C9), we obtain the trivial solution (*α, β, γ*) = (0, 0, 0). Hence, we do not have to consider the case of *t* = 0. Therefore, in the following, we only consider *t* ≠ 0. Replacing constants −*αv*/*t*, −*βv*/*t*, and −*γv*/*t* with *α, β*, and *γ*, we obtain,

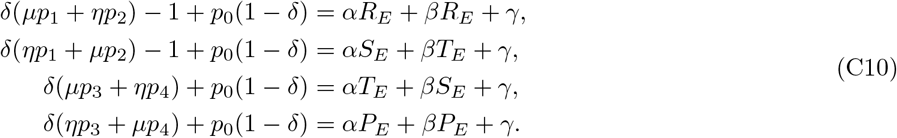

If there exist *α, β*, and *γ* satisfying Eq. (C10), there must be solutions that Eq. (24) holds. This solution corresponds to ZD strategies with observation errors and a discount factor. This is consistent with Eq. (6) in [52] when *δ* =1.

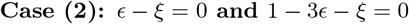

In this case, the equations *ϵ* – *ξ* = 0 and 1 – 3*ϵ* – *ξ* = 0 lead to *ϵ* = 1/4 and *ξ* = 1/4. When *ϵ* =1/4 and *ξ* = 1/4, the expected payoffs *R_E_* = 1/2(*R* + *S*), *S_E_* = 1/2(*R* + *S*), *T_E_* = 1/2(*T* + *P*), and *P_E_* = 1/2(*T* + *P*) hold, which do not satisfy the condition of the prisoner’s dilemma game: *T_E_* > *R_E_* > *P_E_* > *S_E_*. Hence, we can exclude this solution.

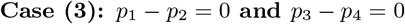

In this case, we substitute *p*_1_ – *p*_2_ = 0 and *p*_3_ – *p*_4_ = 0 into Eq. (C5) to obtain

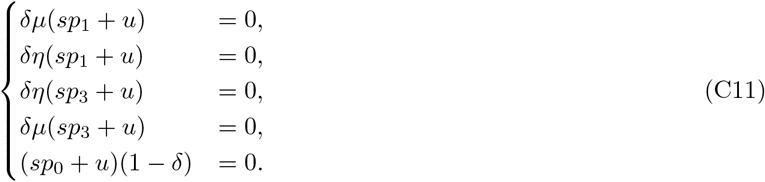

Because the equations *μ* = 0 and *η* = 0 do not hold at the same time and *δ* ≠ 0, we obtain

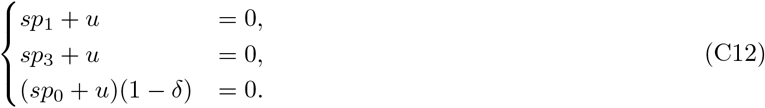

Therefore, we obtain two solutions *p*_0_ = *p*_1_ = *p*_2_ = *p*_3_ = *p*_4_ = −*u/s* or *p*_1_ = *p*_2_ = *p*_3_ = *p*_4_ = −*u/s* and *δ* =1. Both solutions are called unconditional strategies [14, 55]. The former represents unconditional strategies in the case of *δ* ≠ 1. The latter represents unconditional strategies in the case of *δ* = 1. Next, we check whether this solution satisfies Eq. (C4). We substitute the former solution *p*_0_ = *p*_1_ = *p*_2_ = *p*_3_ = *p*_4_ and *u* = −*sp*_0_ into Eq. (C4) to obtain

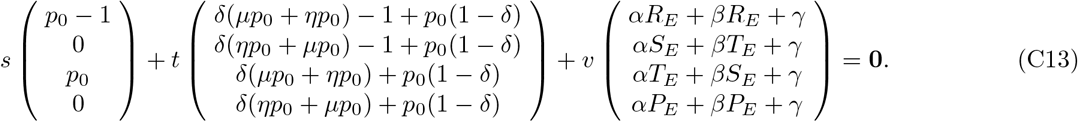

According to *μ* + *η* = 1, we obtain

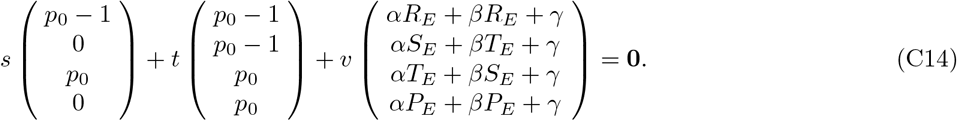

There exist real numbers *s, t, u, v, α, β*, and *γ* which satisfy Eq. (C14) as follows:

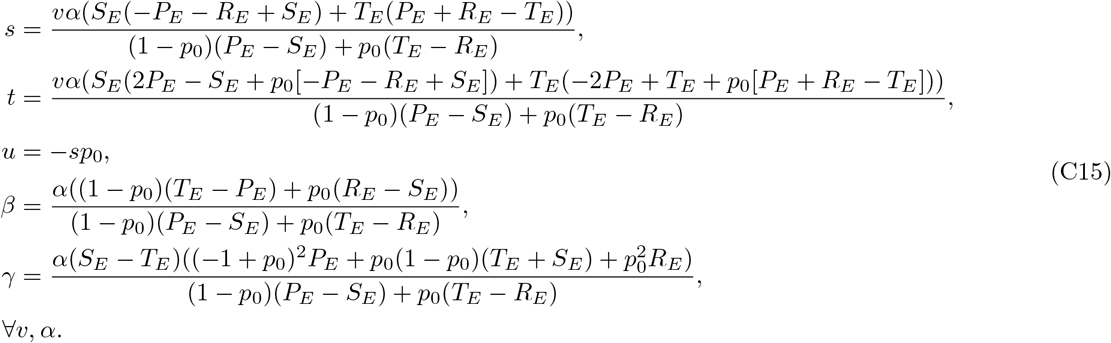

Finally, we substitute the latter solution *p*_1_ = *p*_2_ = *p*_3_ = *p*_4_ = −*u/s* and *δ* = 1 into Eq (C4), we obtain the same real numbers *s, t, u, v, α, β*, and *γ* in the case of the former solution.

This strategy set corresponds to unconditional strategies ***p*** = (*r, r, r, r*; *r*), 0 ≤ *r* ≤ 1. Therefore, the unconditional strategies enforce a linear payoff relationship in the RPD game with both observation errors and a discount factor because there exist real numbers *s, t, u, v, α, β*, and *γ* such that Eqs. (C4) and (C5) are satisfied.

## Appendix D: Calculation of Sec. III.2

We show that neither Extortion nor Generous strategies exist when there are errors. From Eq. (27), we can rewrite ZD strategies as follows:

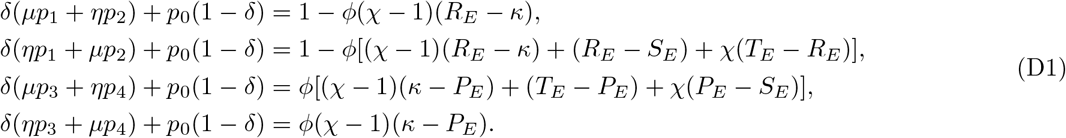

We assume *χ* ≥ 1 because Extortion and Generous are considered. We also assume *η* > 0 because we consider errors. If we solve those equations for *p*_1_, *p*_2_, *p*_3_ and *p*_4_, respectively, we obtain

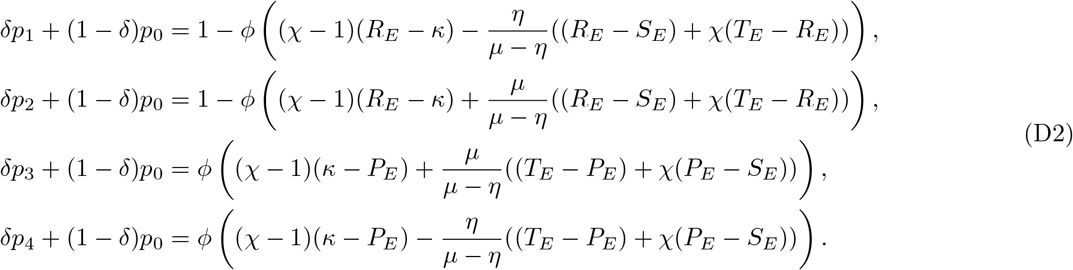

In those equations, when *δ* < 1 and *ϕ* = 0 are given, *p*_1_, *p*_2_, *p*_3_ and *p*_4_ do not exist. When *δ* =1, *ϕ* = 0 is formally allowed, but produces only the singular strategy (1, 1, 0, 0). Thus, we consider the case of *ϕ* ≠ 0.

We substitute *κ* = *P_E_* into the third and fourth equations in Eq. (D2) to obtain

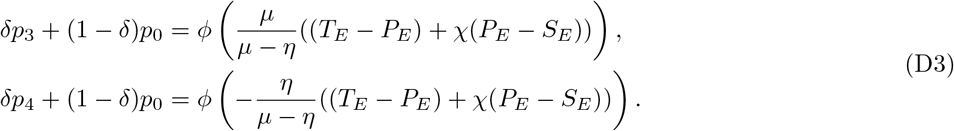

For those two equations, the left sides must be greater than or equal to zero. Thus, the right sides need to be positive due to *ϕ* ≠ 0. When *μ* > *η*, we obtain *ϕ* > 0 from the first equation while *ϕ* < 0 from the second equation. Thus, there is no *ϕ* which satisfies the condition in this case. When *μ* < *η*, we obtain *ϕ* < 0 from the first equation while *ϕ* > 0 from the second equation. Thus, there is also no *ϕ* which satisfies the condition in this case. We exclude the case of *μ* = *η* because this yields *R_E_* = *S_E_* and *T_E_* = *P_E_* which is not the RPD. Therefore, there is no ZD with *κ* = *P_E_*, which corresponds to Extortion, when there are errors.

We substitute *κ* = *R_E_* into the first and second equations in Eq. (D2) to obtain

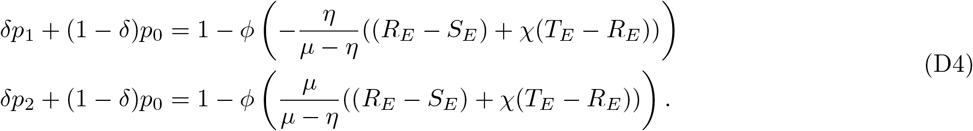

In addition, from 0 ≤ *p_i_* ≤ 1, we obtain

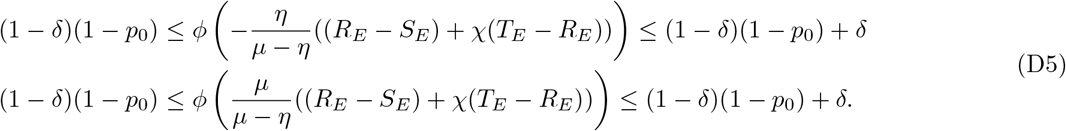

For the inequalities, the middle terms must be positive due to *ϕ* ≠ 0. When *μ* > *η*, we obtain *ϕ* < 0 from the first equation while *ϕ* > 0 from the second equation. Thus, there is no *ϕ* which satisfies the condition in this case. When *μ* < *η*, we obtain *ϕ* > 0 from the first equation while *ϕ* < 0 from the second equation. Thus, there is also no *ϕ* which satisfies the condition in this case. We exclude the case of *μ* = *η* because this yields *R_E_* = *S_E_* and *T_E_* = *P_E_* which is not the RPD. Therefore, there is no ZD with *κ* = *R_E_*, which corresponds to Generous, when there are errors.

## ACKNOWLEDGMENTS

This study was partly supported by the HAYAO NAKAYAMA Foundation for Science & Technology and Culture and JSPS KAKENHI Grants No. JP19K04903 and No. JP19KK0262 (G.I.).

## AUTHOR CONTRIBUTIONS

A.M. and G.I. designed the study and wrote the paper. A.M. conducted mathematical and numerical analyses. G.I. checked the calculation and provided the interpretation of the result.

